# Establishment of Developmental Gene Silencing by Ordered Polycomb Complex Recruitment in Early Zebrafish Embryos

**DOI:** 10.1101/2021.03.16.435592

**Authors:** Graham J.M. Hickey, Candice Wike, Xichen Nie, Yixuan Guo, Mengyao Tan, Patrick Murphy, Bradley R. Cairns

**Affiliations:** Howard Hughes Medical Institute, Department of Oncological Sciences and Huntsman Cancer Institute, University of Utah School of Medicine, Salt Lake City, UT 84112, USA; Department of Biomedical Genetics, Wilmot Cancer Center, University of Rochester School of Medicine, Rochester, NY 14641, USA

## Abstract

Vertebrate embryos achieve developmental competency during zygotic genome activation (ZGA) by establishing chromatin states that silence yet poise developmental genes for subsequent lineage-specific activation. Here, we reveal how developmental gene poising is established *de novo* in preZGA zebrafish embryos. Poising is established at promoters and enhancers that initially contain open/permissive chromatin with ‘Placeholder’ nucleosomes (bearing H2A.Z, H3K4me1, and H3K27ac), and DNA hypomethylation. Silencing is initiated by the recruitment of Polycomb Repressive Complex 1 (PRC1), and H2Aub1 deposition by catalytic Rnf2 during preZGA and ZGA stages. During postZGA, H2Aub1 enables Aebp2-containing PRC2 recruitment and H3K27me3 deposition. Notably, preventing H2Aub1 (via Rnf2 inhibition) eliminates recruitment of Aebp2-PRC2 and H3K27me3, and elicits transcriptional upregulation of certain developmental genes during ZGA. However, upregulation is independent of H3K27me3 – establishing H2Aub1 as the critical silencing modification at ZGA. Taken together, we reveal the logic and mechanism for establishing poised/silent developmental genes in early vertebrate embryos.

**Impact Statement:** *De novo* polycomb domains are formed in zebrafish early embryos by focal histone H2Aub1 deposition by Rnf2-PRC1 – which imposes transcription silencing – followed by subsequent recruitment of Aebp2-PRC2 and H3K27me3 deposition.

## Introduction

Early vertebrate embryos initiate embryonic/zygotic transcription, termed zygotic genome activation (ZGA), and must distinguish active housekeeping genes from developmental genes, which must be temporarily silenced, but kept available for future activation (Chan et al., 2019; Lindeman et al., 2011; Murphy et al., 2018; Potok et al., 2013; Vastenhouw et al., 2010).

Developmental gene promoters in early embryos are packaged in ‘active/open’ chromatin – which can involve a combination of histone variants (e.g. H2A.Z), open/accessible chromatin (via ATAC-seq), permissive histone modifications (e.g. H3K4me1/2/3, H3K27ac), and (in vertebrates) focal DNA hypomethylation (Akkers et al., 2009; Bogdanovic et al., 2012; Chan et al., 2019; Jiang et al., 2013; Lindeman et al., 2011; Murphy et al., 2018; Potok et al., 2013; Vastenhouw et al., 2010). As H3K9me3 and H3K27me3 are very low or absent at ZGA, it remains unknown how developmental gene silencing occurs at ZGA within an apparently permissive chromatin landscape, and how subsequent H3K27me3 is established at developmental genes during postZGA stages.

We addressed these issues in zebrafish, which conduct full ZGA at the tenth synchronous cell cycle of cleavage stage (∼3.5 hours post fertilization (hpf), ∼2000 cells)(Schulz et al., 2019). Prior to ZGA (preZGA), zebrafish package the promoters and enhancers of housekeeping genes and many developmental genes with chromatin bearing the histone variant H2afv (a close orthologue of mammalian H2A.Z, hereafter termed H2A.Z(FV)), and the ‘permissive’ modifications H3K4me1 and H3K27ac (Murphy et al., 2018; Zhang et al., 2018); a combination termed ‘Placeholder’ nucleosomes – as they hold the place where poising/silencing is later imposed (Murphy et al., 2018). Curiously, at ZGA in zebrafish, (and also in mice and humans), silent developmental gene promoters also contain H3K4me3, a mark that normally resides at active genes (Dahl et al., 2016; Lindeman et al., 2011; Liu et al., 2016; Vastenhouw et al., 2010; Xia et al., 2019; Zhang et al., 2016). After ZGA, developmental genes progressively acquire H3K27me3 via deposition by Polycomb Repressive Complex 2 (PRC2) (Lindeman et al., 2011; Liu et al., 2016; Vastenhouw et al., 2010; Xia et al., 2019). Therefore, the central issues are how developmental genes bearing Placeholder nucleosomes and H3K4me3 are transcriptionally silenced during preZGA and ZGA stages in the absence of H3K27me3, and how subsequent H3K27me3 is focally established during postZGA (Figure 1 - figure supplement 1A).

**Figure 1:**
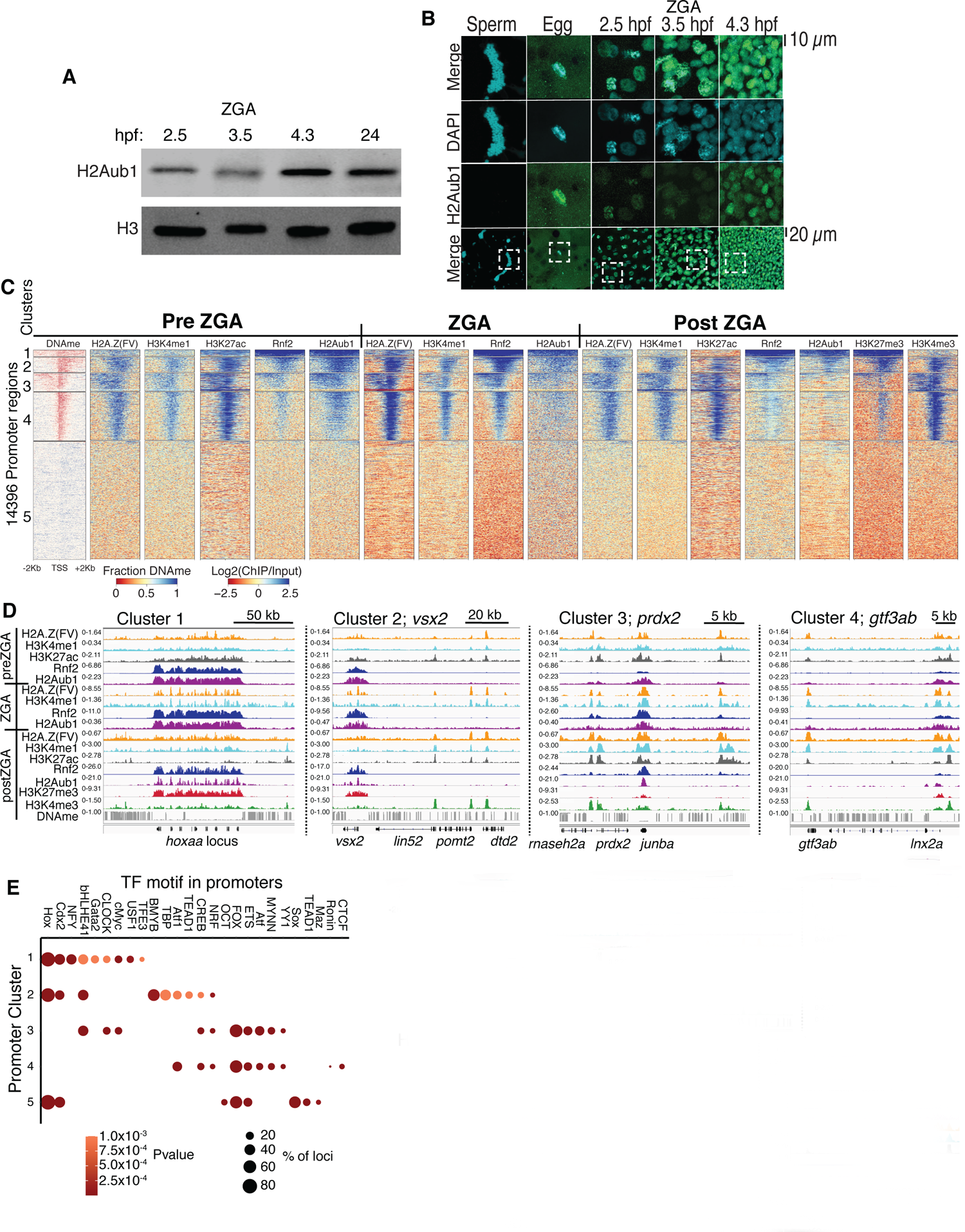
PRC1 occupancy and activity precedes H3K27me3 establishment at promoters. **(A)** Detection of H2Aub1 and histone H3 (control) in chromatin extracts by western blot, prior to (2.5 hpf), during (3.5 hpf), following (4.3hpf) ZGA, and 24 hpf. **(B)** Nuclear H2Aub1 immunofluoresence in zebrafish sperm, oocytes (egg), and embryos prior to (2.5 hpf), during (3.5 hpf), and following (4.3hpf) ZGA. Bottom row: the dashed square indicates the field of view in upper panels. **(C)** K-means clustering of whole-genome bisulfite sequencing (WGBS, for DNAme) and ChIP-seq (histone modifications/variant) at promoters (UCSC refseq). DNAme heatmap displays WGBS fraction methylated scores (note: red color indicates regions that lack DNAme). ChIP-seq heatmaps display log2(ChIP/input) scores. **(D)** Genome browser screenshots of ChIP-seq enrichment at representative genes from the indicated K-means clusters in panel (C). **(E)** Transcription factor motif enrichment from HOMER (Heinz et al., 2010) at promoter clusters from (C).

**Figure 1 – figure supplement 1:**
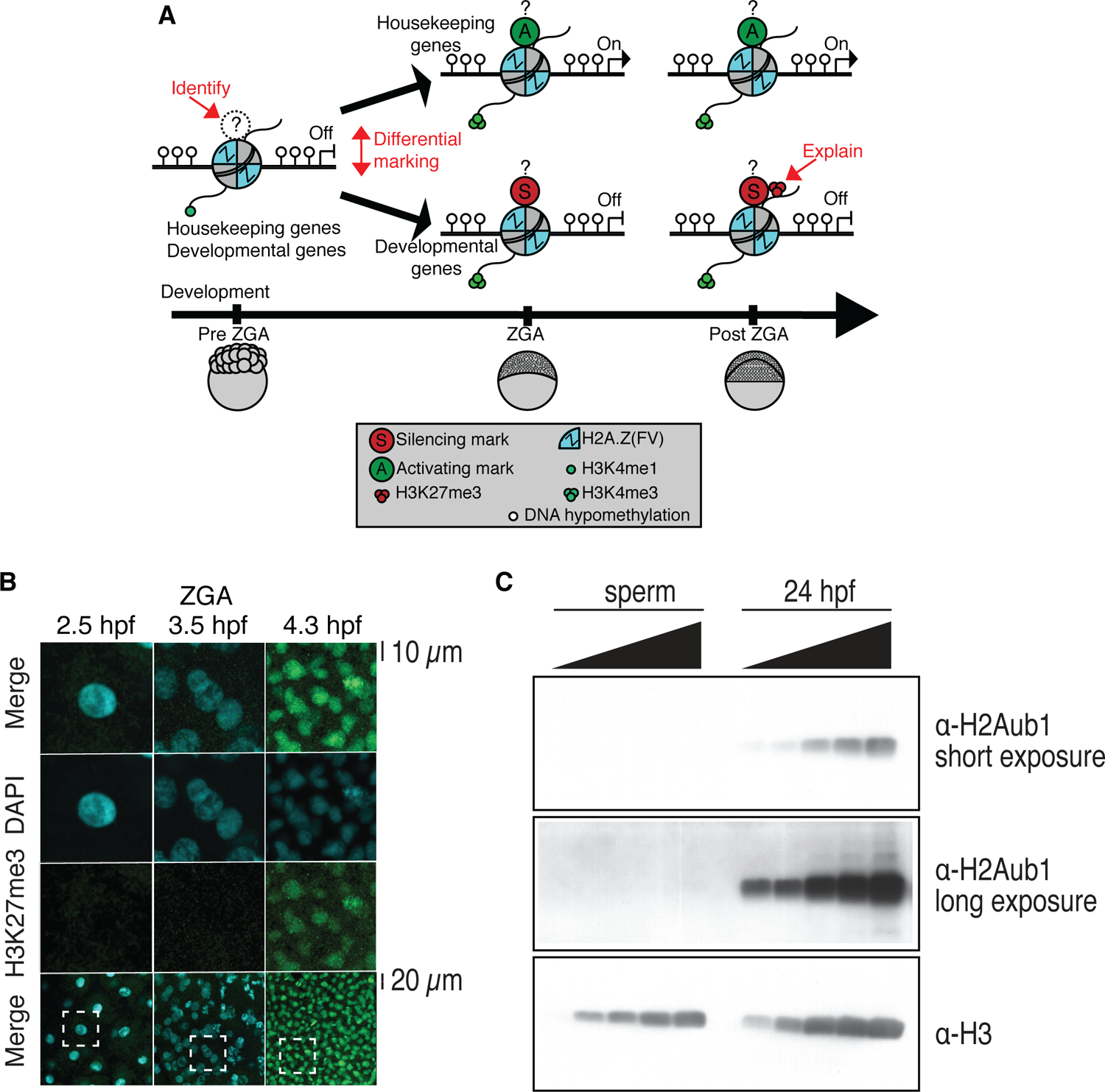
Profiling of histone marks in zebrafish embryos and sperm. **(A)** A model illustrating the current knowledge gaps regarding how developmental gene silencing and housekeeping gene activation is achieved in early vertebrate embryos. Question marks represent the possible existence of histone modifications, activating (A?) or silencing (S?) that distinguish gene classes prior to zygotic genome activation (ZGA). **(B)** Nuclear H3K27me3 detection by immunofluorescence in zebrafish embryos following ZGA (4.3hpf). 1 of 3 biological replicates is shown. No H3K27me3 staining was observed in embryos at preZGA (2.5hpf) or ZGA (3.5 hpf). Bottom row: the dashed square indicates denotes the field of view for the corresponding top 3 rows. **(C)** H2Aub1 is undetected by western blot in zebrafish sperm. Varying amounts of protein from zebrafish sperm (left) or 24hpf embryos (right) were used for western blotting. Top: short exposure probing for H2Aub1. Middle: long exposure probing for H2Aub1. Bottom: histone H3 loading control. 1 of 3 biological replicates is shown.

**Figure 1 – figure supplement 2:**
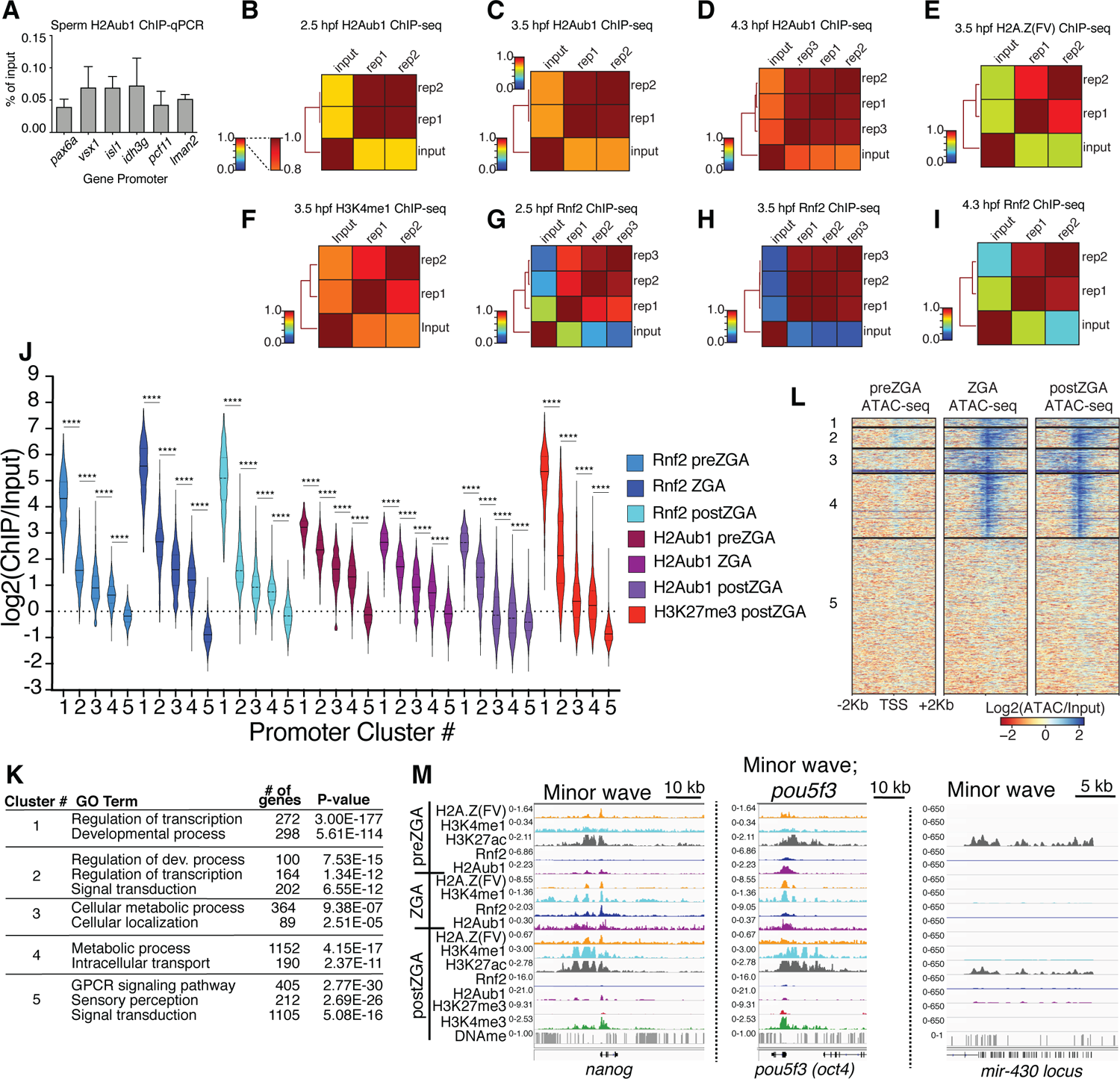
ChIP replicate structures. **(A)** ChIP-qPCR of H2Aub1 of indicated gene promoters in zebrafish sperm. No H2Aub1 enrichment was detected at developmental gene promoters (pax6a, vsx1, isl1) in sperm. Housekeeping gene promoters (idh3g, pcf11, lman2) were used as negative control regions. Three biological replicates were conducted. **(B-I)** Genome-wide correlation matrices of indicated ChIP-seq replicates and input from zebrafish embryos at the indicated developmental timepoints. Heatmap matrices display Pearson correlation coefficients (B-D, G-I) or Spearman correlation coefficients (E, F). **(J)** Violin plots of ChIP-seq enrichment levels at promoter K-means clusters displayed in Figure 1C. A 1Kb window flanking the TSS was utilized to collect mean log2(ChIP/Input) data. Unpaired t-test with Welch’s correction was performed in GraphPad Prism version 8.3.1. Asterisks represent P values that are less than 0.0001. **(K)** GO-term analysis of promoter clusters in Figure 1C. Top GO terms (enriched gene categories) for the promoter clusters are shown, along with the number of genes (#) and the P-value for enrichment. GO-term analysis was performed with DAVID (Huang da et al., 2009). **(L)** Heatmaps of ATAC-seq at UCSC RefSeq promoters. K-means clusters from Figure 1C were used to plot data. **(M)** Genome browser screenshots of representative genes involved in the minor wave of zygotic transcription with ChIP-seq enrichment from clusters in Figure 1C.

## Results

### H2Aub1 is Present in PreZGA Zebrafish Embryo Chromatin

First, we sought a repressive histone modification that might explain how developmental genes are silenced at ZGA. We examined zebrafish embryos at preZGA (2.5 hpf, ∼256 cells), ZGA (3.5 hpf ∼2000 cells) and postZGA (4.3 hpf, >4K cells) – and confirmed very low-absent H3K27me3 (Figure 1 - figure supplement 1B) (Lindeman et al., 2011; Vastenhouw et al., 2010) and H3K9me3 absence (Laue et al., 2019) during preZGA and ZGA, but revealed the presence of histone H2A monoubiquitination at lysine 119 (termed hereafter H2Aub1), a repressive mark deposited by the Polycomb Repressive Complex 1 (PRC1) (Figure 1A, B, one of three biological replicates is displayed) (de Napoles et al., 2004; Kuroda et al., 2020; Wang et al., 2004).

Notably, zebrafish sperm lacked H2Aub1 whereas oocytes displayed H2Aub1 (Figure 1B; Figure 1 - figure supplement 1C; Figure 1 - figure supplement 2A). Current antibodies were not designed to distinguish H2A.Z(FV)ub1 from H2Aub1, so hereafter we refer to the epitope as H2Aub1.

### Developmental Promoters Acquire Placeholder, Rnf2, and H2Aub1 During PreZGA

To localize H2Aub1 we conducted chromatin immunoprecipitation (ChIP) experiments at preZGA, ZGA and postZGA (replicate structures for Figure 1 - figure supplement 2B-D), and examined promoters (Figure 1) and enhancers (Figure 2). For all ChIP experiments, 2-3 biological replicates were conducted, which involved isolating different batches of zebrafish embryos. To more finely examine ZGA, we conducted additional ChIP profiling of Placeholder nucleosomes at ZGA (3.5 hpf), which complemented our prior profiling at preZGA (2.5 hpf) and postZGA (4.3 hpf) (Murphy et al., 2018) (replicate structure for Figure 1- figure supplement 2E, F). Interestingly, we found H2Aub1 highly co-localized at gene promoters and enhancers with Placeholder nucleosomes, H3K27ac (Zhang et al., 2018) (Figure 1C), and ATAC-seq sensitive chromatin (Figure 1 - figure supplement 2L, two biological replicates) during preZGA and ZGA stages. However, during postZGA, high levels of H2Aub1 overlap with only a portion of Placeholder-bound loci, an observation explored further, below.

**Figure 2:**
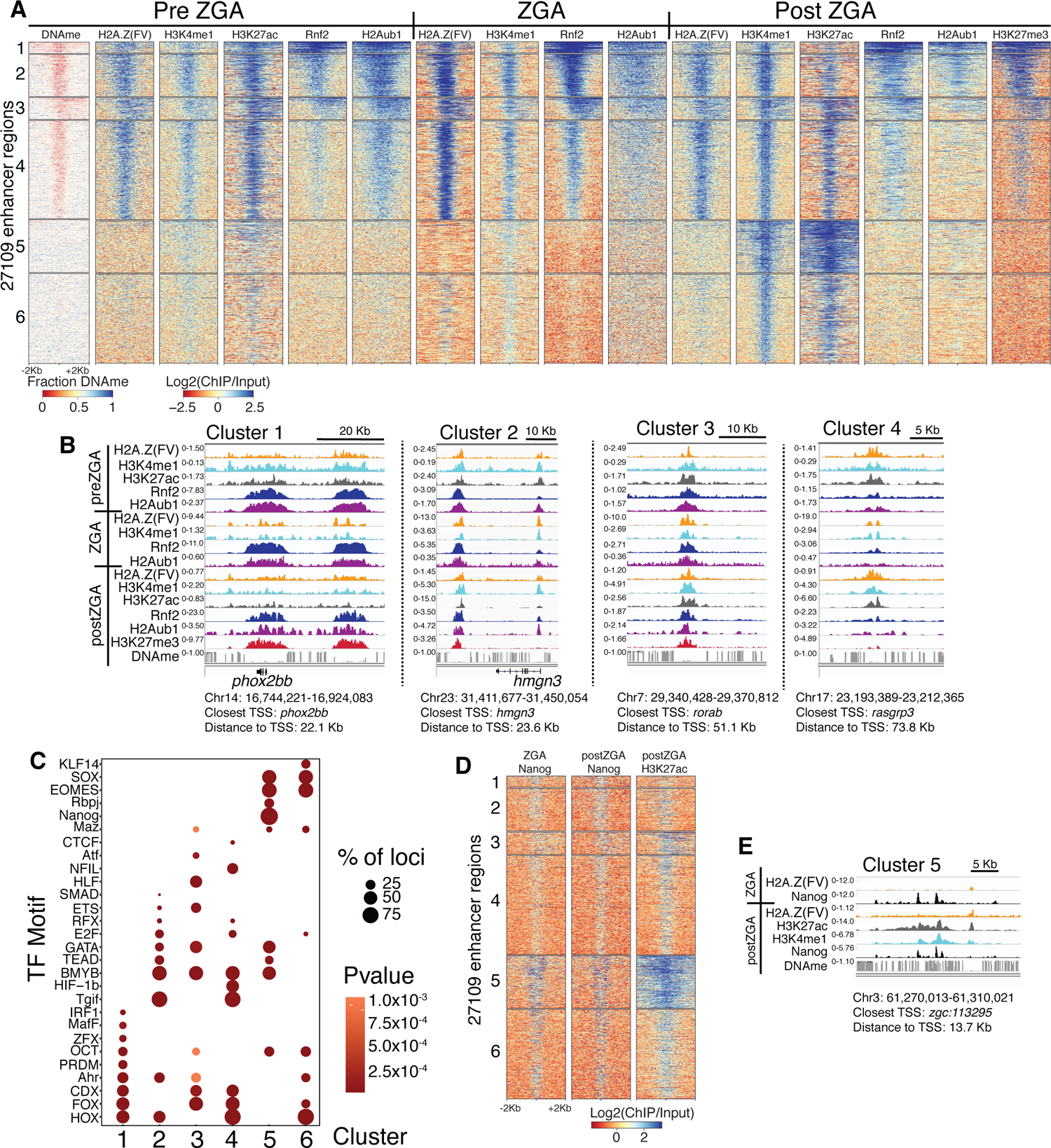
PRC1 occupancy and activity precedes H3K27me3 at enhancers. **(A)** K-means clustering of WGBS (for DNAme) and ChIP-seq at enhancers (postZGA H3K4me1 peak summits located outside of promoters). DNAme heatmap displays WGBS fraction methylated scores (note: red color indicates regions that lack DNAme). ChIP-seq heatmaps display log2(ChIP/input) scores. **(B)** Genome browser screenshots of ChIP-seq enrichment at representative loci from the indicated K-means clusters in (A). **(C)** Transcription factor motif enrichment from HOMER (Heinz et al., 2010) at enhancer clusters from (A). **(D)** Features of an enhancer cluster with exceptionally high H3K27ac and Nanog binding. K-means clusters generated in (A) were utilized to plot heatmaps of Nanog and H3K27ac. **(E)** A genome browser screenshot depicting Nanog and H3K27ac ChIP enrichment at a DNA-methylated enhancer from cluster 5 in (D).

**Figure 2 – figure supplement 1:**
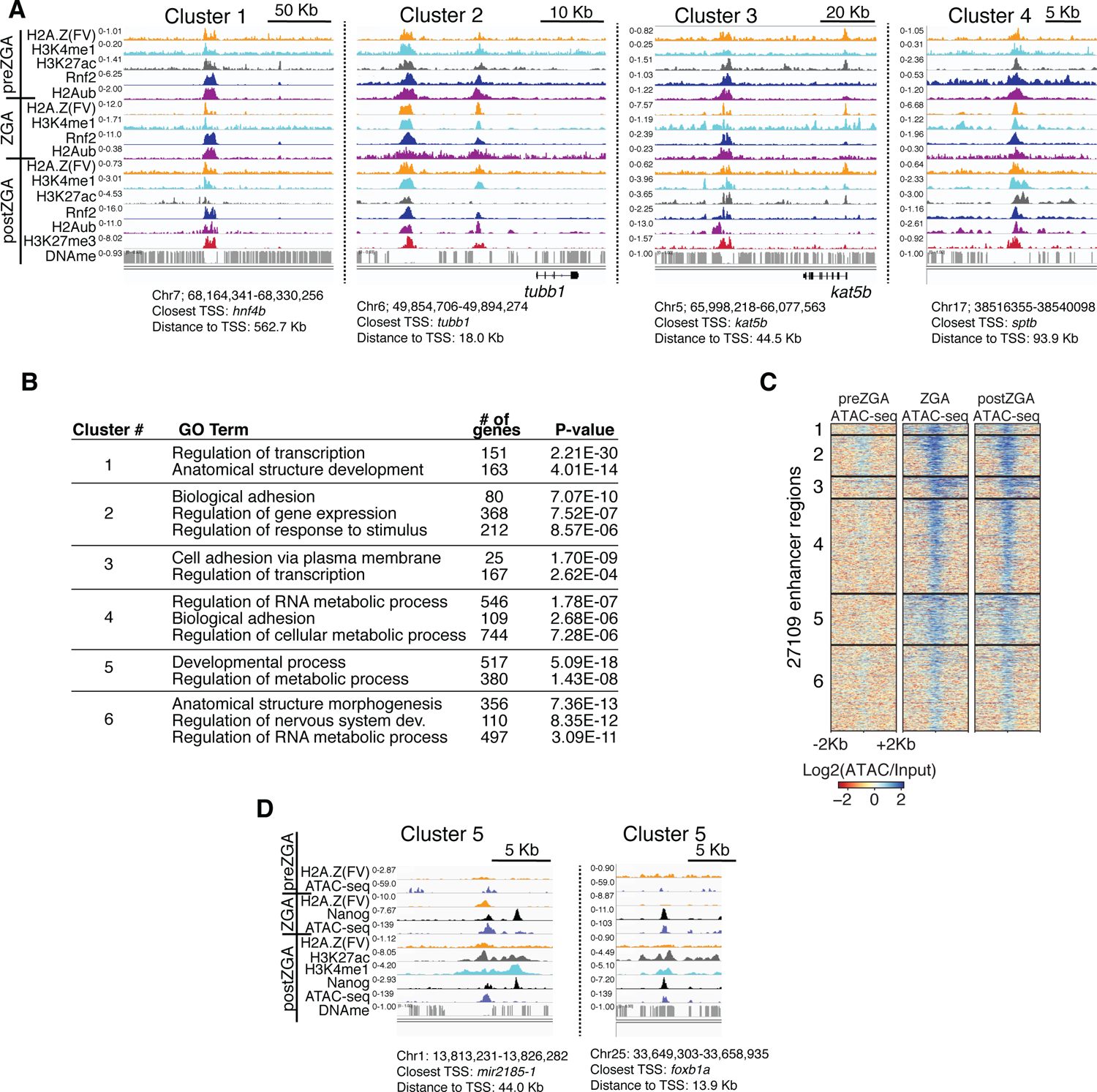
Additional examples of ChIP enrichment at enhancers. **(A)** Genome browser screenshots of ChIP-seq enrichment at representative enhancer loci from enhancer K-means clusters in Figure 2A. **(B)** GO-term analysis of enhancer clusters in Figure 1C. Top GO terms (enriched gene categories) for the nearest genes to the enhancer clusters are shown, along with the number of genes (#) and the P-value for enrichment. GO-term analysis was performed with DAVID (Huang da et al., 2009). **(C)** Heatmaps of ATAC-seq at H3K4me1 peak summits. K-means clusters from Figure 2A were used to plot data. **(D)** Genome browser screenshots of representative ATAC-seq and ChIP-seq enrichment at enhancer cluster 5 from Figure 2A & 2D.

Rnf2 is the sole zebrafish ortholog of Ring1a and Ring1b/Rnf2, the mutually-exclusive E3 ligases within mammalian PRC1, which adds monoubiquitin (ub1) to H2A and H2A.Z (de Napoles et al., 2004; Le Faou et al., 2011; Wang et al., 2004). Rnf2 ChIP-seq at preZGA, ZGA and postZGA revealed striking co-incidence with H2Aub1, and clustering by Rnf2 occupancy revealed five promoter chromatin types, which differed in Rnf2, H2Aub1, and H3K27me3 enrichment, and gene ontology (Figure 1C; Figure 1 - figure supplement 2G-K). Regions with high H2Aub1 and Rnf2 involve broad clustered loci encoding developmental transcription factors (cluster 1, e.g *hoxaa*) or narrow solo/dispersed developmental transcription factors (cluster 2, e.g. *vsx2*) (Figure 1C, D; GO analysis for Figure 1 - figure supplement 2K). In counter distinction, loci bearing Placeholder and low-moderate levels of H2Aub1 and low Rnf2 largely constitute housekeeping/metabolic genes, with either broad (cluster 3, e.g. *prdx2*) or narrow (cluster 4, e.g. *gtf3ab*) H3K4me3 and H3K27ac occupancy at postZGA (Figure 1C, D; Figure 1 – figure supplement 2K). Notably, ‘minor wave’ ZGA genes (genes transcribed at 2.5hr), including those for pluripotency (e.g. *nanog, pou5f3/oct4*), bear marking similar to housekeeping genes, and the robustly-transcribed *mir430* locus appears markedly enriched in H3K27ac at preZGA (Figure 1 - figure supplement 2M) (Chan et al., 2019). Finally, cluster 5 promoters contain DNAme, and lack Placeholder, H2Aub1 and Rnf2. Thus, over the course of ZGA, loci with Placeholder nucleosomes resolve into two broad classes of loci: developmental genes with high PRC1 and H2Aub1 (clusters 1&2), and housekeeping (or ‘minor wave’) genes that lack substantial PRC1 and H2Aub1, but contain high H3K4me3 and H3K27ac at postZGA (clusters 3&4) (Figure 1C, D; Figure 1 - figure supplement 2J, K).

### H3K27me3 Establishment Occurs During PostZGA, and Only at Locations Pre-marked with High Rnf2 and H2Aub1

We find robust H3K27me3 deposition occurring during postZGA at promoters marked during preZGA with high Rnf2 and H2Aub1, specifically at clusters 1 and 2 (Figure 1C, D).

Furthermore, as embryos transition from preZGA to postZGA, H3K27ac diminishes at developmental loci, whereas housekeeping genes (clusters 3 and 4 Figure 1C, D) retain strong H3K27ac and become active. During postZGA, developmental genes acquire the combination of low-moderate H3K4me3 and high H3K27me3, termed ‘bivalency’ (Azuara et al., 2006; Bernstein et al., 2006). Interestingly, promoters that become bivalent postZGA involve those pre-marked with higher relative Rnf2 and H2Aub1, whereas promoters with high H3K27ac, high H3K4me3 and low-absent H3K27me3 postZGA involve those pre-marked with lower relative Rnf2 and H2Aub1 (Figure 1C, D; Figure 1 - figure supplement 2J). This observation raised the possibility that high H2Aub1 levels may help recruit PRC2 to subsequently deposit H3K27me3.

To identify candidate transcription factors (TFs) that might bind selectively at the promoters of particular clusters, we analyzed the DNA sequences flanking the TSS (500bp) at each cluster using the motif finding program, HOMER (Figure 1E) (Heinz et al., 2010). Largely non-overlapping motifs were identified for TF binding sites at clusters linked to developmental versus housekeeping genes (Figure 1E, clusters 1&2 versus 3&4; partitioned by H2Aub1/Rnf2 levels). Here, the strong enrichment of motifs for homeodomain-containing TFs (and other families) in clusters 1 and 2 provides candidate factors that may help recruit PRC1 to developmental loci. DNA methylated promoters at ZGA (cluster 5) represent a large and heterogeneous set of genes, which are activated in particular cell types later in development.

### Enhancer Poising Parallels Features and Factors at Promoters

Analysis of enhancer regions revealed features that were similar to those at developmental promoters (Figure 2). Specifically, enhancers with high H2Aub1 and Rnf2 during preZGA and ZGA acquired robust H3K27me3 during post ZGA (Figure 2A, B; Figure 2 – figure supplement 1A). Enhancers with low/absent Rnf2 and H2Aub1 failed to attract robust H3K27me3 at postZGA, instead bearing high levels of H3K27 acetylation (Figure 2A, B; Figure 2 - figure supplement 1A). Candidate TF binding sites were enriched at enhancers (Figure 2C), and these sites overlapped partly with those enriched at promoters, consistent with the expectation that the factors that recruit histone modifiers to promoters and enhancers partially overlap. Taken together, enhancers acquire H3K27me3 during postZGA in proportion to their levels of Rnf2 and H2Aub1 during preZGA and ZGA, consistent with our observations at promoters.

### A Unique Enhancer Class with High H3K27ac and DNA Methylation

Curiously, enhancer cluster 5 (Figure 2A) was unique – displaying high H3K4me1, very high H3K27ac, and open chromatin (via ATAC-seq analysis; Figure 2 - figure supplement C, D) – but bore DNA methylation – an unusual combination given the typical strong correlation between high H3K4me1 and DNA hypomethylation. Notably, Nanog binding sites were highly enriched solely at cluster 5, and Nanog ChIP-seq during ZGA and postZGA (Xu et al., 2012) showed Nanog occupancy highly and selectively enriched at cluster 5 relative to other enhancer clusters (Figure 2C-E). Thus, cluster 5 enhancers may utilize Nanog and H3K27ac to open and poise these DNA methylated enhancers for later/subsequent transition to an active state. Consistent with this notion, GO analysis of cluster 5 enriches for terms related to developmental and signaling processes (p-value; 5.1E-18) (Figure 2 - figure supplement 1B).

### The PRC2 Component Aebp2 is Co-incident with Rnf2 and H2Aub1 at Developmental Loci

The pre-marking of developmental genes with H2Aub1 and Rnf2-PRC1 prior to H3K27me3 establishment raised the possibility of a ‘non-canonical’ (nc) mode of recruitment, involving ncPRC1 action (H2Aub1 addition) followed by the recruitment of PRC2 (ncPRC2) – via H2A/Zub1 recognition – to deposit H3K27me3. This mode and order of recruitment has precedent in *Drosophila* and in mammalian ES cell cultures, with the Aebp2 and Jarid2 protein components of ncPRC2 recognizing H2Aub1 and both targeting and facilitating H3K27me3 addition (Blackledge et al., 2014; Blackledge et al., 2020; Cooper et al., 2014; Cooper et al., 2016; Kalb et al., 2014; Kasinath et al., 2021; Tamburri et al., 2020). We then addressed whether establishment of H3K27me3 during postZGA is mediated by the non-canonical Aebp2-PRC2 complex at loci pre-marked with H2Aub1. Interestingly, Aebp2 protein levels were very low during preZGA and ZGA stages, but robustly detected postZGA (Figure 3A), without a large increase in *aebp2* transcript levels (Figure 3 - figure supplement 1A). Aebp2 ChIP-seq during postZGA revealed a remarkably high coincidence of Aebp2 with H2Aub1, Rnf2 and H3K27me3 at promoters (Figure 3B, C; Figure 3 - figure supplement 1B-D) and at enhancers (Figure 3D, E, Figure 3 - figure supplement 1E). Taken together, these results suggest that translational upregulation and/or protein stability enables Aebp2 protein accumulation postZGA – enabling the ‘reading/binding’ of H2Aub1, and H3K27me3 deposition during postZGA by PRC2.

**Figure 3:**
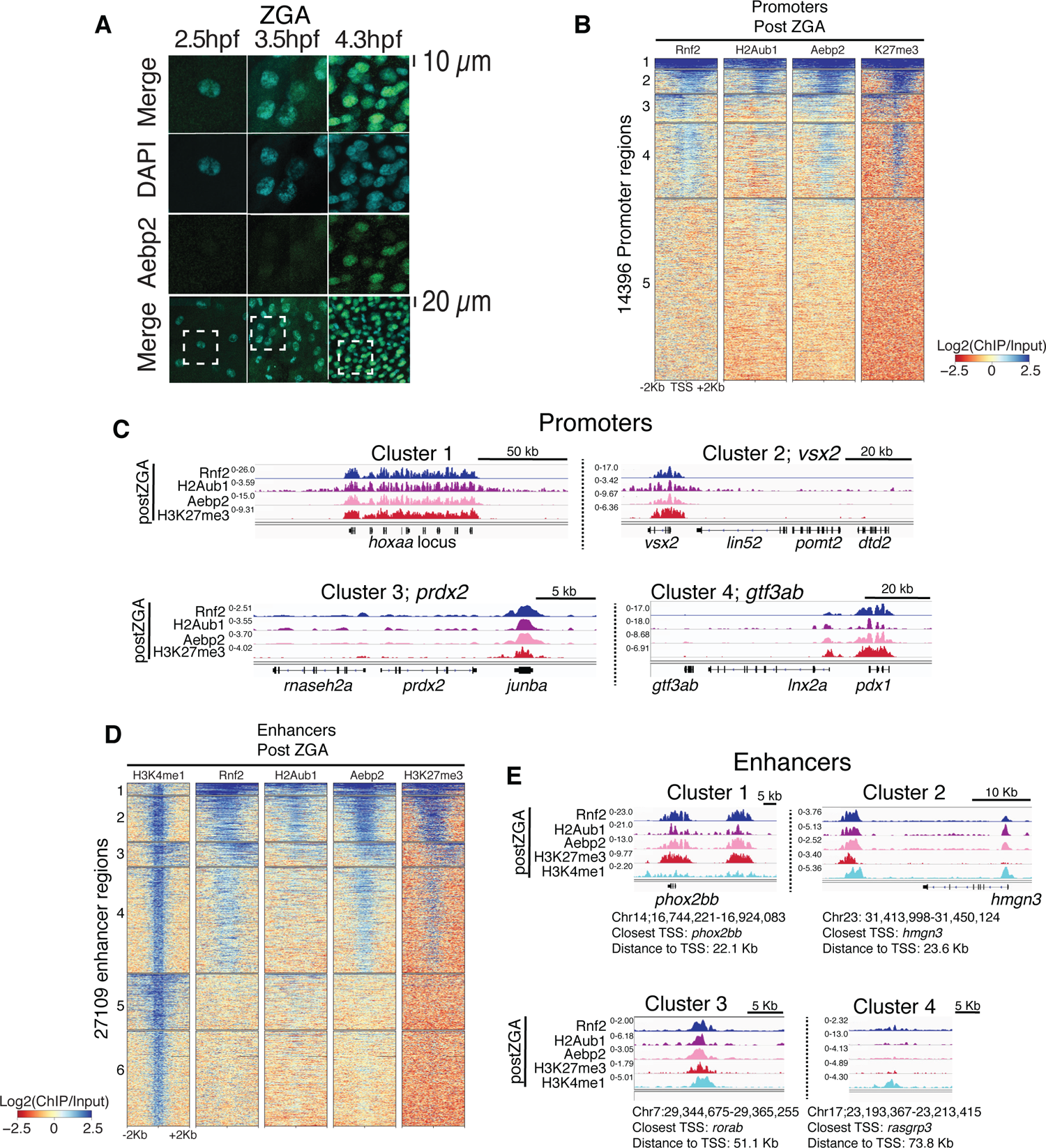
Aebp2-PRC2 mediates *de novo* H3K27me3 at loci pre-marked by H2Aub1. **(A)** Nuclear Aebp2 detection by immunofluoresence in postZGA zebrafish embryos (4.3 hpf). No Aebp2 staining was detected at preZGA (2.5 hpf) or ZGA (3.5 hpf). Bottom row: the dashed square indicates the field of view in upper panels. 1 of 3 biological replicates is shown. **(B)** Aebp2 binding at promoters during postZGA overlaps and scales with occupancy of Rnf2, H2Aub1, and H3K27me3. Promoter clusters from Figure 1C were utilized to plot heatmaps. **(C)** Genome browser screenshots of ChIP-seq at representative promoter loci from clusters in (B). **(D)** Aebp2 binding at enhancers during postZGA overlaps with occupancy of Rnf2, H2Aub1, and H3K27me3. Enhancer clusters from Figure 2A were utilized to plot heatmaps. **(E)** Genome browser screenshots of ChIP-seq enrichment at representative enhancer loci from clusters in (D).

**Figure 3 – figure supplement 1:**
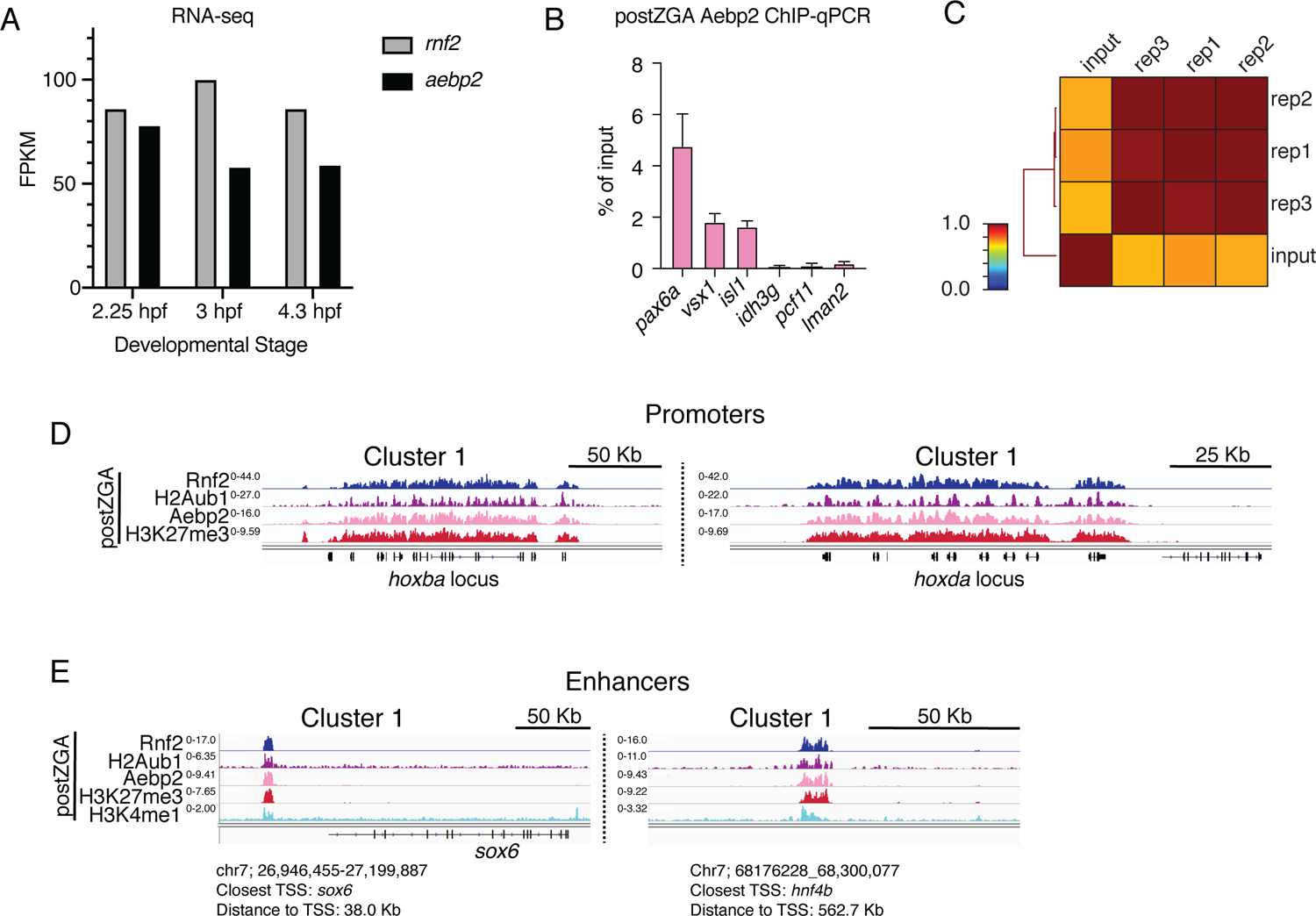
ChIP profiling of Aebp2 in postZGA embryos. **(A)** Abundance of *aebp2* mRNA in early zebrafish embryos measured by RNA-sequencing. Increased Aebp2 protein levels from preZGA to postZGA observed in Figure 4A are not explained by increased aebp2 mRNA levels (black bars). Levels of *rnf2* mRNA are plotted for comparison (grey bars). RNA-seq data was collected from http://www.ebi.ac.uk/gxa/ex-periments/E-ERAD-475. **(B)** Aebp2 ChIP-qPCR demonstrates enrichment of Aebp2 at developmental gene promoters (*pax6a, vsx1, isl1*) in postZGA stage embryos (4.3 hpf). Housekeeping gene promoters (*idh3g, pcf11, lman2*) were used as negative control regions. Note: promoter regions assayed here are either robustly marked by H2Aub1 (*pax6a, vsx1, isl1*) or lack H2Aub1 (*idh3g, pcf11, lman2*). Three biological replicates were conducted for ChIP. For qPCR, three technical replicates were conducted on each ChIP biological replicate. **(C)** Genome wide correlation matrix of Aebp2 ChIP-seq replicates and input from postZGA (4.3 hpf) zebrafish embryos. Heatmap matrix displays Pearson correlation coefficients. **(D)** Genome browser screenshots of representative promoters with ChIP-seq enrichment demonstrating binding of Aebp2 at promoters bearing Rnf2, H2Aub1, and H3K27me3 at post ZGA (4.3 hpf). **(E)** Genome browser screenshots of representative enhancers with ChIP-seq enrichment demonstrating binding of Aebp2 at enhancers bearing Rnf2, H2Aub1, and H3K27me3 at post ZGA (4.3 hpf).

### Loss of H2Aub1 via Rnf2 Inhibition Prevents Aebp2 Localization and H3K27me3 Deposition

To functionally test whether H2Aub1 recruits Aebp2-PRC2 for *de novo* establishment of H3K27me3 at developmental genes, we utilized the RNF2 inhibitor, PRT4165 (Chagraoui et al., 2018; Ismail, McDonald, Strickfaden, Xu, & Hendzel, 2013; Zhu et al., 2018). PRT4165 is a small molecule inhibitor of Rnf2 that has previously been shown to strongly reduce H2Aub1 modification, but not to affect the activity of related H2A E3 ligases such as Rnf8 and Rnf168 (Chagraoui et al., 2018; Ismail et al., 2013; Zhu et al., 2018). In each of three biological replicates, PRT4165 treatment (150µM) from the 1-cell stage onward largely eliminated H2Aub1 by 4hpf (ZGA) (Figure 4A, B), and conferred a developmental arrest that resembled untreated 4 hpf embryos (Figure 4 - figure supplement 1A). To determine whether the loss of H2Aub1 conferred loss of Aebp2 genomic targeting we performed ChIP experiments on Aebp2 in PRT4165-treated and DMSO-treated embryos (3 biological replicates per condition).

**Figure 4:**
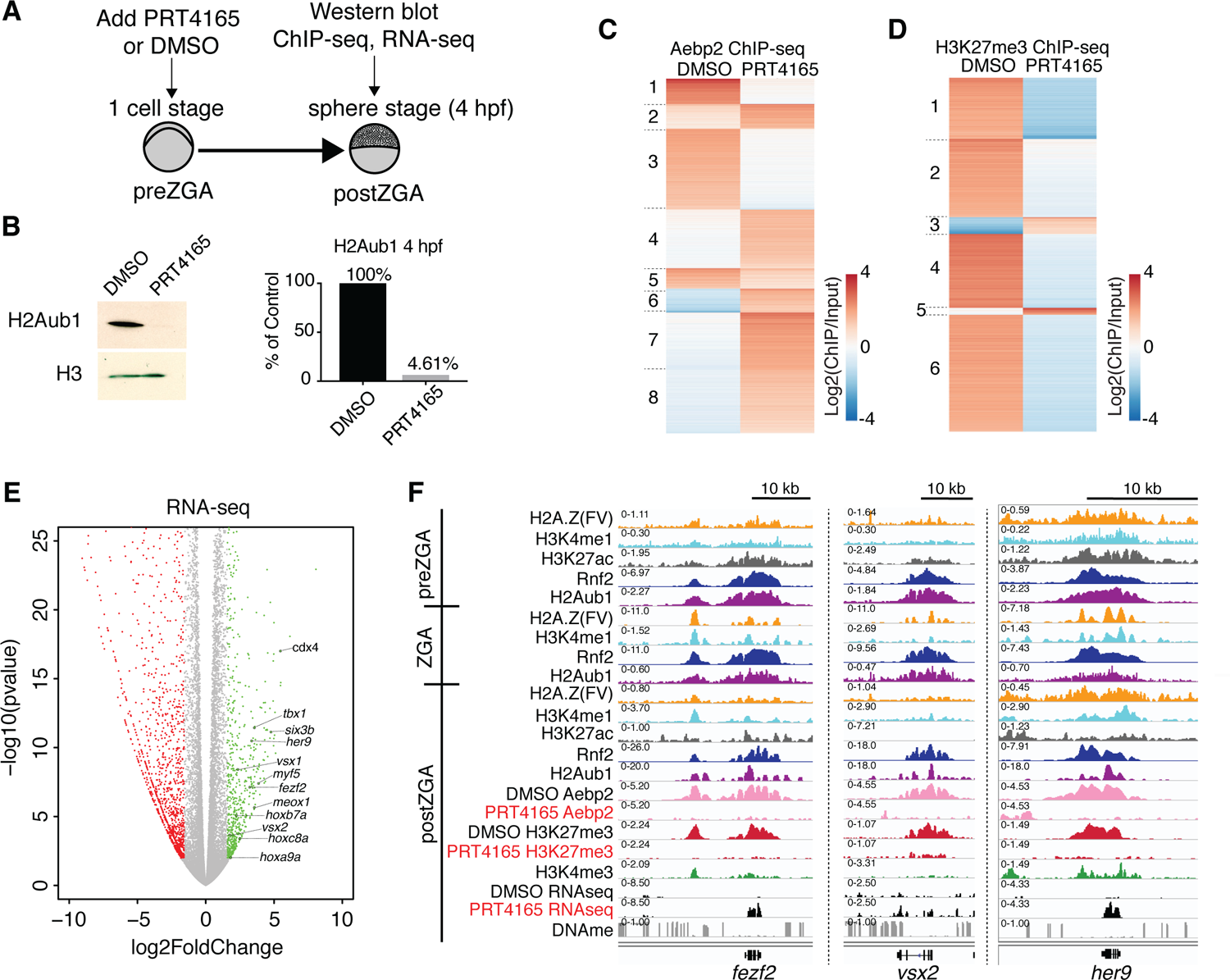
Catalytic activity of PRC1 is required for aebp2 binding, H3K27me3 establishment, and transcriptional repression of developmental genes. **(A)** Experimental design of drug treatments to inhibit Rnf2 activity. Embryos at the 1-cell stage were added to media containing either PRT4165 (150 μM) or DMSO and raised until 4 hpf. (B) PRT4165 treatment of embryos confers bulk loss of H2Aub1 at 4hpf. Left: western blot for H2Aub1 in 4hpf embryos treated with DMSO (vehicle) or 150 μM PRT4165 (Rnf2 inhibitor). Right: quantification western blot in left panel. (C) Impact of H2Aub1 removal on Aebp2 genomic localization. K-means clustering of Aebp2 ChIP-seq enrichment at all loci with called peaks in embryos (4 hpf) treated from the 1-cell stage with either DMSO or 150 μ M PRT4165. (D) Impact of H2Aub1 removal on H3K27me3 genomic localization. K-means clustering of Aebp2 ChIP-seq enrichment at all loci with called peaks in embryos (4 hpf) treated from the 1-cell stage with either DMSO or 150 μM PRT4165. (E) Impact of Rnf2 inhibition on gene expression.

Remarkably, developmental loci that normally display high Aebp2 in untreated or DMSO-treated embryos lost Aebp2 binding following PRT4165 treatment (Figure 4C, clusters 1, 3, & 5; Figure 4 - figure supplement 1B, C). Curiously, PRT4165 treatment also conferred limited new/ectopic Aebp2 peaks (Figure 4C, clusters 4, 6, 7,8), however our profiling of H3K27me3 following PRT4165 treatment (3 biological replicates) revealed that new/ectopic Aebp2 sites did not acquire H3K27me3 (Figure 4 - figure supplement 2B). Importantly, treatment with PRT4165 eliminated or strongly reduced H3K27me3 at virtually all loci normally occupied by H3K27me3 (Figure 4D, Figure 4 - figure supplement 1D), a conclusion supported by our use of a ‘spike-in’ control involving *Drosophila* nuclei bearing H3K27me3-marked regions in all 3 replicates per condition. Curiously, a small number of ectopic H3K27me3 peaks were observed in the PRT4165 treatment, however these loci were not bound by Aebp2 (Figure 4 - figure supplement 2B). Taken together, H2Aub1 deposition by Rnf2 during preZGA is required for the recruitment of Aebp2 and subsequent *de novo* deposition of H3K27me3 postZGA at developmental loci.

**Figure 4 – figure supplement 1:**
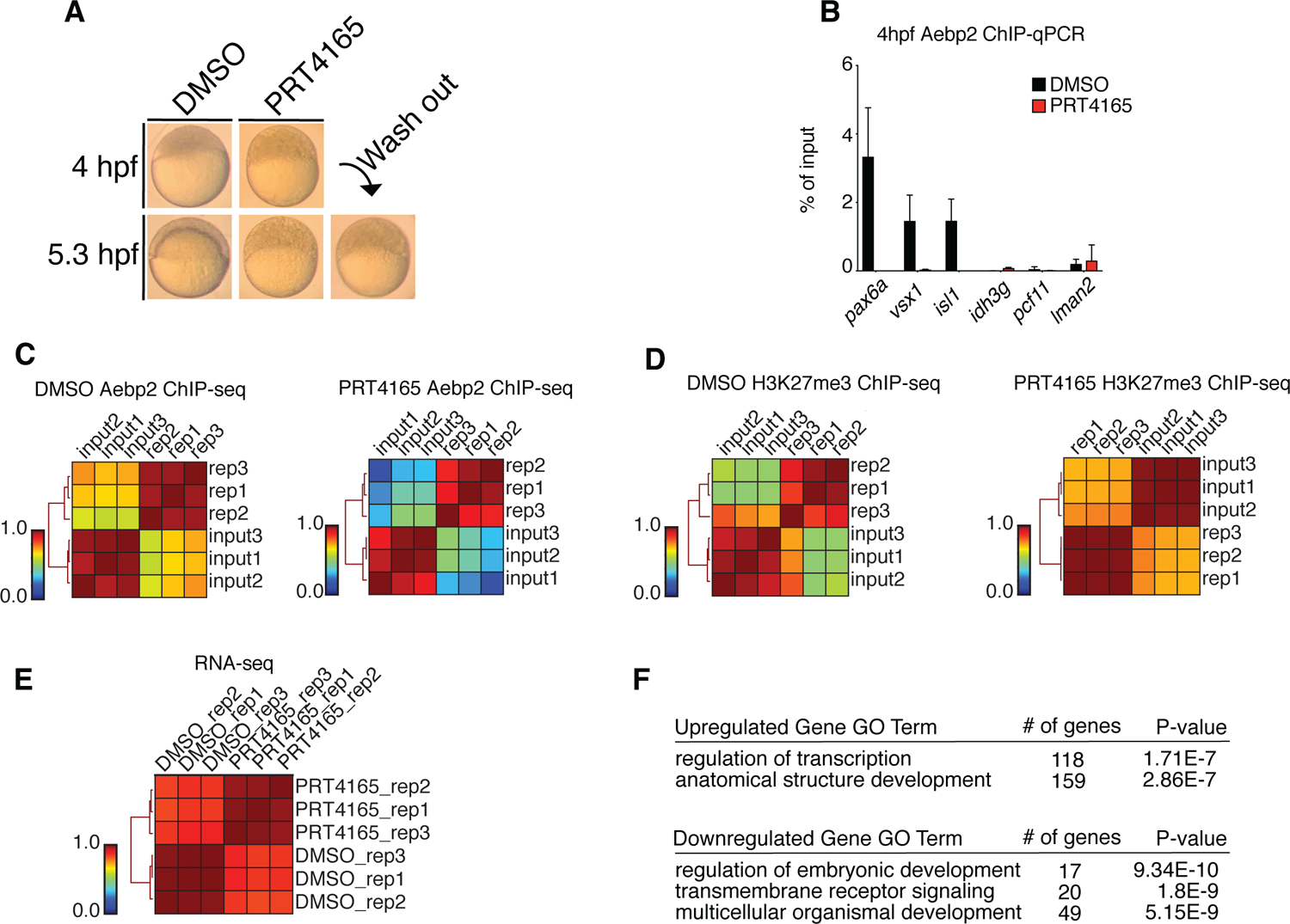
Replicate structures of ChIP-seq and RNA-seq experiments involving drug treatments. **(A)** Treatment of zebrafish embryos with PRT4165 (150 μM) beginning at the 1 cell stage results in developmental arrest at 4hpf. Removal of PRT4165 at 4 hpf failed to restore normal developmental progression to embryos (lower right, “Wash out”). **(B)** Aebp2 ChIP-qPCR in postZGA (4 hpf) embryos treated with DMSO (black bars) or PRT4165 (150 μM, red bars). Binding of Aebp2 to developmental gene promoters (*pax6a, vsx1, isl1*) is lost upon PRC1 inhibition. **(C)** Genome wide correlation matrices of Aebp2 ChIP-seq replicates and input from postZGA (4 hpf) zebrafish embryos treated with DMSO (left) or PRT4165 (150 μM, right). Heatmap matrices display Pearson correlation coefficients. **(D)** Genome wide correlation matrices of H3K27me3 ChIP-seq replicates and input from postZGA (4 hpf) zebrafish embryos treated with DMSO or PRT4165 (150 μM, right). Heatmap matrices display Pearson correlation coefficients. **(E)** Genome wide correlation matrix of RNA-seq replicates from postZGA (4 hpf) zebrafish embryos treated with DMSO or PRT4165. Heatmap matrix displays Pearson correlation coefficients. **(F)** GO-terms associated with upregulated & downregulated transcripts depicted in figure 4E. GO-term analysis was performed with DAVID (Huang da et al., 2009).

**Figure 4 – figure supplement 2:**
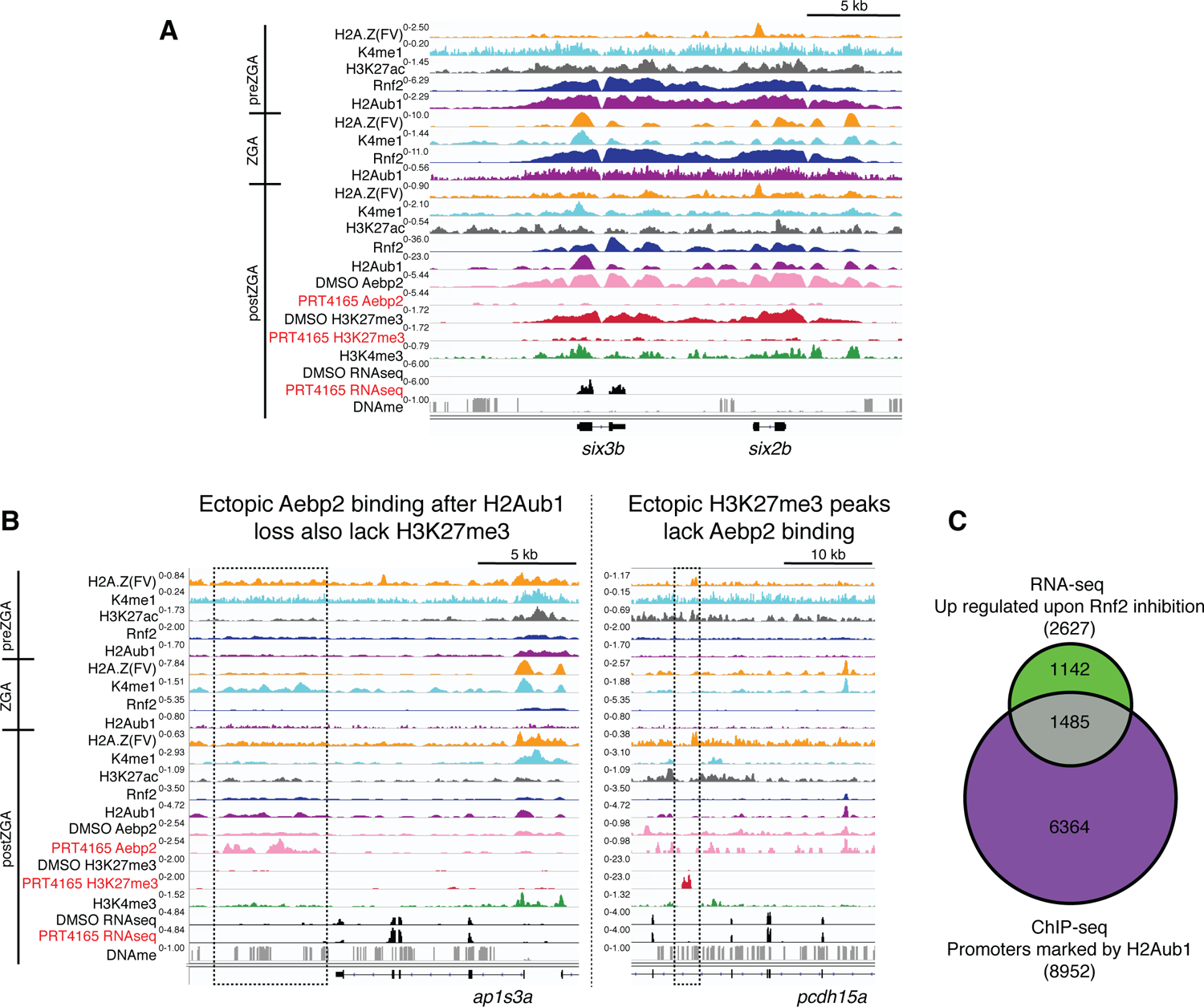
Loss of H2Aub1 results in disrupted Aebp2 localization, H3K27me3 marking and transcriptional de-repression. **(A)** Genome browser screenshots of representative de-repressed genes upon PRC1 inhibition. Both six3b and six2b are bound by Rnf2 and marked by H2Aub1 from preZGA onward. At postZGA these genes normally attract binding by Aebp2 and acquire H3K27me3. Inhibition of Rnf2 (via PRT4165) causes failed Aebp2 binding, and prevention of H3K27me3 establishment. Notably, six3b is precociously expressed upon loss of H2Aub1. **(B)** Genome browser screenshots of representative loci that gain ectopic Aebp2 binding (left) or H3K27me3 marking (right) upon PRC1 inhibition. Ectopic binding events for Aebp2 or H3K27me3 appear to be mutually exclusive. Dotted boxes denote region of ectopic binding or marking by Aebp2 or H3K27me3, respectively. **(C)** Approximately 16.6% of genes normally silenced by H2Aub1 promoter marking are transcriptionally upregulated upon inhibition of Rnf2. Venn diagram displays the overlap between the gene promoters normally marked by H2Aub1 and protein coding genes that are transcriptionally upregulated (p-value <0.001, >1.5 fold change) after Rnf2 inhibiton. Here H2Aub1 marked promoters are defined as regions +/-2Kb from transcription start sites and having peak(s) called by MACS2 (Feng et al., 2012) (q-value cutoff 0.01) of H2Aub1 from 4.3 hpf H2Aub1 ChIP-seq.

To determine whether H2Aub1 impacts transcriptional repression of developmental genes at ZGA, we performed RNA sequencing (3 biological replicates per condition) on 4hpf embryos that were either vehicle-treated (DMSO) or PRT4165-treated from the 1-cell stage onward (Figure 4 - figure supplement 1E), and identified both up- and downregulated genes (Figure 4E). PRT4165-upregulated and downregulated genes (>3-fold, p-value <0.01) were both enriched in developmental factors, but the number of genes associated with upregulated GO-terms were substantially greater than downregulated GO-terms (Figure 4 - figure supplement 1F). Here, ∼16.6% of H2Aub1-marked protein coding genes were upregulated, which may reflect the availability at ZGA of an opportunistic activator, following H2Aub1 loss (Figure 4 - figure supplement 2C). Affected genes include those in clustered loci (e.g. *Hox* genes) where the effect was moderate, as well as non-clustered/solo formats where the effect of PRT4165 was more pronounced (Figure 4E, F; Figure 4 - figure supplement 2A). Taken together, these results strongly suggest that H2Aub1 represses developmental genes during preZGA and ZGA, independent of H3K27me3, and that the absence of H2Aub1 renders developmental genes susceptible to precocious activation following ZGA, which confers developmental arrest prior to gastrulation.

Volcano plot of RNA-seq data from PRT4165-treated versus untreated embryos (4hpf). Green and Red data points signify transcripts with P-values <0.01 and at least a 3-fold change (increase or decrease) in expression, respectively. Marquee upregulated genes encoding developmental transcription factors are labelled. **(F)** Genome browser screenshots of representative developmental genes which, upon Rnf2 inhibition, lose Aebp2 binding and H3K27me3 marking, and become transcriptionally active.

## Discussion

Our work reveals that developmental gene silencing in early zebrafish embryos is established through sequential recruitment and activity of PRC1 and PRC2 complexes, respectively, to otherwise open/permissive loci bearing Placeholder nucleosomes (Figure 5). Placeholder nucleosomes contain H3K4me1 and the histone variant H2A.Z(FV), are installed by the chromatin remodeler SRCAP during preZGA (Murphy et al., 2018), and are focally pruned to small regions by the chaperone Anp32e (Kobor et al., 2004; Krogan et al., 2003; Mao et al., 2014; Obri et al., 2014; Wang et al., 2016). Here, our reanalysis of published data confirms that H3K27ac is an additional component of Placeholder nucleosomes during preZGA (Zhang et al., 2018). Functional studies reveal that Placeholder nucleosomes prevent DNAme where they are installed, and are utilized to reprogram the DNAme patterns during cleavage stage. Therefore, from a ‘permissive’ Placeholder platform during preZGA, two very different chromatin/transcriptional states are attained at ZGA: active or poised (Murphy et al., 2018).

**Figure 5:**
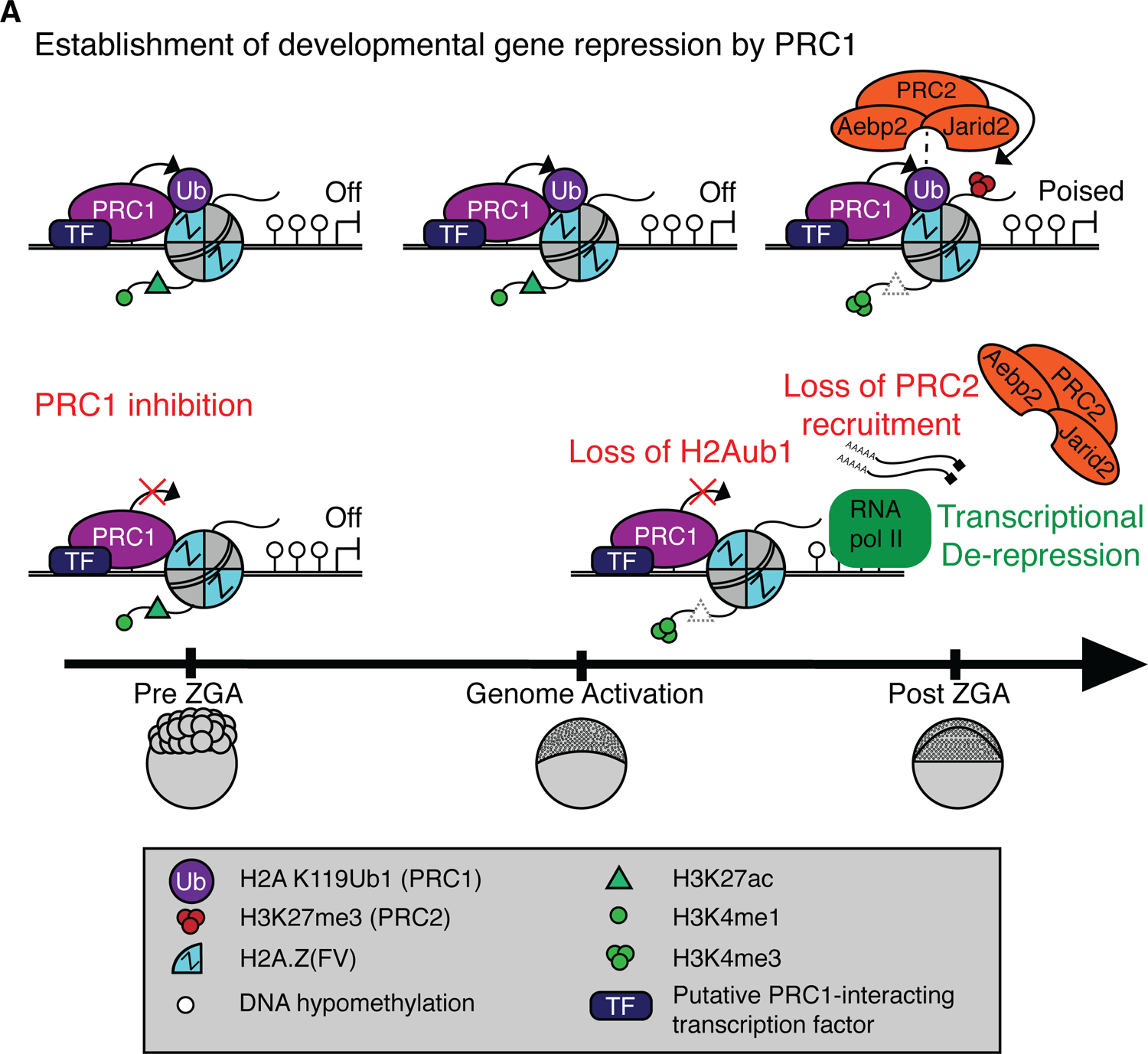
Model for Polycomb-mediated establishment of developmental gene silencing during zebrafish embryogenesis. **(A)** Prior to ZGA, Rnf2-PRC1 is recruited by transcription factors (TF) to promoters (shown) and enhancers (not shown) of developmental genes bearing Placeholder nucleosomes (H2A.Z(FV), H3K4me1, H3K27ac). Rnf2-PRC1 deposition of H2Aub1 recruits Aebp2-PRC2 to catalyze H3K27me3 addition. H2Aub1 ablation (via Rnf2 inhibition) eliminates Aebp2-PRC2 recruitment and prevents H3K27me3 establishment. Notably, H2Aub1 loss causes precocious transcription of certain developmental genes after ZGA, identifying H2Aub1 as a critical component of silencing at ZGA.

Our work suggests that the initial poised state at developmental genes and enhancers involves the imposition of polycomb-based silencing upon the permissive states established by Placeholder nucleosomes. Specifically, we observe H2Aub1 addition by Rnf2/PRC1 during preZGA and ZGA to confer initial transcriptional silencing at developmental loci – which is subsequently read by the Aebp2/PRC2 complex to add H3K27me3 after ZGA (Figure 5). Prior work in ES cells has provided an *in vitro* parallel in which PRC1 activity (H2Aub1 addition) can precede PRC2 recruitment at certain loci (Blackledge et al., 2014; Blackledge et al., 2020; Cooper et al., 2014; Cooper et al., 2016; Kalb et al., 2014; Tamburri et al., 2020; Tavares et al., 2012), which has been termed ‘non-canonical’ order of recruitment, to contrast with prior data showing the reverse/canonical order (Wang et al., 2004). Furthermore, and consistent with our work, human AEBP2 and JARID2 have recently have been shown to directly bind H2Aub1 and stimulate PRC2 activity in the presence of H3K4 methylation (Kasinath et al., 2021), and a similar mechanism may be utilized to establish bivalency after ZGA in zebrafish.

Our work also clarifies and extends prior work in zebrafish which showed that an incross of zebrafish heterozygous for an *rnf2* loss-of-function truncation mutation yielded a pleitropic terminal phenotype at 3 days post fertilization (dpf), coincident with the loss of *rnf2* RNA. Notably, loss of *rnf2* at 3 dpf was associated with partial upregulation of certain developmental genes, but H3K27me3 remained fully present, showing that Rnf2 is not required for H3K27me3 maintenance (Chrispijn et al., 2019). However, progression of *rnf2* mutants to 3 dpf may have relied on maternally-inherited WT *rnf2* RNA or protein to provide the initial establishment of gene silencing and H3K27me3, raising the possibility that *rnf2* is actually essential at a much earlier developmental stage. Our use of the Rnf2 inhibitor, PRT4165, during preZGA and ZGA stages reveals the necessity for Rnf2 activity for the establishment of developmental gene silencing, for subsequent H3K27me3 addition, and for progression of zebrafish development beyond the ZGA stage. Importantly, developmental gene upregulation is not is attributable to H3K27me3 loss, as maternal zygotic *ezh2* mutant zebrafish embryos do not precociously activate developmental genes during ZGA, and they progress through gastrulation without H3K27me3 (Rougeot et al., 2019; San et al., 2016; San et al., 2019). Notably, although upregulation of developmental genes in the presence of PRT4165 is clear, this involves ∼16.6% of the developmental gene repertoire occupied by H2Aub1/Rnf2. Here, we suggest that tissue-specific activators for the majority of developmental genes are not present at ZGA.

One curiosity arising from our work is why zebrafish utilize the non-canonical PRC1 complex, which adds monoubiquitination to H2A/H2A.Z(FV), rather than canonical PRC1 complex, which compacts chromatin for initial developmental gene silencing. Here, we speculate that the rapid (∼16 minute) cell cycles that characterize the preZGA cleavage state – coupled to the need for continual DNA replication during cleavage stage – are not compatible with the compaction conferred by canonical PRC1 (Gao et al., 2012; Grau et al., 2011; Lau et al., 2017). Furthermore, the necessary substrate to recruit canonical PRC1 to chromatin, H3K27me3, is absent in preZGA embryos. Instead, the use of the repression modes conducted by non-canonical PRC1 addition of H2Aub1 – which antagonizes RNA Pol II transcriptional initiation or bursting, may help confer silencing without conferring a compaction that might impede DNA replication (Dobrinić et al., 2020; Stock et al., 2007). However, once embryos exit cleavage stage, the cell cycle greatly lengthens, and Aebp2-PRC2 complexes add H3K27me3 to loci – which may then enable canonical PRC1 to localize to and conduct compaction. Future studies are necessary to address this possible temporal order and role for canonical PRC complexes.

Here, we observe that developmental or housekeeping gene promoters attract either a high or low level PRC1 binding, respectively. How ncPRC1 is recruited to CpG islands in ES cells is partially understood, as the ncPRC1 subunit Kdm2b helps recruit ncPRC1 to CpG islands of developmental genes. In this context, Kdm2b binds to hypomethylated CpGs via the CxxC motif, (Blackledge et al., 2014; Farcas et al., 2012; He et al., 2013; Wu et al., 2013). However, ncPRC1 is recruited robustly to only a minority of Kdm2b-bound CpG islands, implying that additional factors are needed to specify recruitment of ncPRC1 to CpG islands of developmental genes.

Here, we speculate that particular transcription factors, such as the candidates in Figures 1E and 2C, are likewise utilized in cooperation with zebrafish Kdm2b to enable strong focal recruitment of Rnf2-PRC1 to Placeholder-occupied developmental loci. In contrast, the candidate transcription factors at housekeeping genes would recruit MLL complexes to implement H3K4 methylation, and not Rnf2-PRC1. Finally, we expect our work here will motivate work to explore whether mammals likewise initially utilize Ring1a/Rnf2-associated PRC1 complexes to add H2Aub1 to help establish developmental gene poising and regulation, and specify particular regions for subsequent marking by H3K27me3.

## Materials and Methods

### Zebrafish Husbandry

Wild type Tübingen zebrafish were maintained as described (Westerfield, 2007). All experiments involving zebrafish were approved by University of Utah IACUC (Protocol 20-04011). Embryos were scored for developmental staged as described (Kimmel et al., 1995).

### Acid Extraction of Nuclear Proteins

200 embryos were added to 1.5 ml tubes and washed twice with cold PBS. 800 µl of Mild Cell Lysis Buffer (10 mM Tris-HCl pH 8.1, 10 mM NaCl, 0.5 NP-40, 2X protease inhibitors) was applied to embryos and incubated on ice for 5 minutes. Embryos were homogenized by passing through a 20 gauge syringe several times. Samples were briefly centrifuged to bring down chorions. Supernatants were transferred to fresh tubes and centrifuged at 1300 X g for 5 minutes at 4°C. Pelleted nuclei were washed twice with cold Mild Cell Lysis Buffer. Nuclei were resuspended to a final volume of 800 µl in cold Mild Cell Lysis Buffer and supplemented with 10 µl of sulfuric acid (18.4M). Samples were sonicated for 10 seconds (1 second on, 0.9 seconds off) at 30% output using a Branson sonicator. Proteins were extracted for 30 minutes at 4°C on a rotator. 160 µl of 100% trichloroacetic acid was added and proteins were allowed to precipitate for 30 minutes on ice. Samples were centrifuged at 13000 rpm for 5 minutes at 4°C. Protein pellets were washed with 800 µl of cold acidified acetone, and centrifuged again at 13000 rpm for 5 minutes at 4°C. Protein pellets were washed with 800 µl of cold acetone, and centrifuged again at 13000 rpm for 5 minutes at 4°C. Supernatant was discarded and pellets were dried at 37°C for 5 minutes. Dried protein pellets were resuspended in 2X Laemli sample buffer and boiled for 8 minutes. Samples were then used for western blotting. Bands were quantified in Image J (Schneider et al., 2012).

### Immunohistochemistry & DAPI staining

Standard protocol for immunohistochemistry was followed as described (Fernández et al., 2013; Zhang et al., 2018). Three biological replicates were performed for each immunohistochemistry experiment. Briefly, 30 embryos were collected at appropriate time points and fixed with fresh 4% paraformaldehyde (Electron Microscopy, Cat # 50980487) in 1xPBS at room temperature for 12 hours. Droplets of glacial acetic (100%, Merck, Cat # 1000560001) or DMSO (final concentration 0.5%, Sigma) were added 5–10 sec after initiation of the fixation. Chorions were manually removed from fixed embryos with forceps and dechorionated embryos were dehydrated in methanol and stored at −20°C. For immune-staining, embryos were rehydrated into PB3T (1xPBS with 0.3% TritonX-100, and then incubated in blocking agent (1% BSA, 0.3 M glycine in PB3T). Embryos were incubated with primary antibodies diluted in blocking agent overnight at 4°C. Primary antibodies were removed and embryos were washed extensively with PB3T. Embryos were next incubated with appropriate secondary antibodies in the dark followed by extensive washes in PB3T. Primary antibodies used for immune-staining are listed below.

Secondary antibodies used were donkey α-rabbit IgG-488 at 1:500 (Life Technologies, Cat # A-21206). DAPI was used at 1:1000 as a nuclear counterstain. The yolk cells were removed from embryo and embryo was mounted on glass slide with ProLong Gold Antifade mounting media (Thermo Fisher, Cat# P-36931) and a 2.0mm square coverslip sealed with nail polish. Samples were stored at 4°C until imaged.

### Imagining of Zebrafish Embryos

Images were acquired on a Leica SP8 White Light laser confocal microscope. Image processing was completed using Nikon NIS-Elements multi-platform acquisition software with a 40X/1.10 Water objective. Fiji (ImageJ, V 2.0.0-rc-69/1.52p) was utilized to color DAPI channel to cyan, GFP color remained green. Confocal images are max projections of Z stacks taken 0.5µm apart for a total of the embryo ∼7-12 µm.

### Primary Antibodies

The following antibodies were utilized in for the present study: anti-H2Aub1 (Cell Signaling Technology Cat# 8240; RRID:AB_10891618), anti-Rnf2 (Cell Signaling Technology Cat# 5694; RRID:AB_10705604), anti-H2A.Z (Active Motif Cat# 39113; RRID:AB_2615081), anti-H3K4me1 (Active Motif Cat# 39297; RRID:AB_2615075), anti-Aebp2 (Cell Signaling Technology Cat# 14129; RRID: AB_2798398), anti-H3K27me3 (Active Motif Cat# 39155; RRID: AB_2561020), anti-H3 (Active Motif Cat# 39763; RRID: AB_2650522).

### ChIP-Seq in Zebrafish Embryos

#### Embryo Fixation

Approximately 1.5 million cells were used for each ChIP replicate. Embryos were allowed to progress to the desired developmental stage and then transferred to 1.5 ml microcentrifuge tubes (∼200 embryos per tube). Chorions were removed enzymatically by treatment with pronase 1.25mg/ml in PBS). Dechorionated embryos were gently washed twice with PBS to remove pronase. Samples were fixed with 1% formaldehyde (Electron Microscopy Sciences, Cat# 15712) for 10 minutes at room temperature with end over end rotation. Fixation was quenched with 130 mM glycine for five minutes at room temperature. Samples were centrifuged for five minutes at 500 x g at 4°C. Supernatant was discarded and cell pellets were washed twice with ice cold PBS. Cell pellets were frozen with liquid nitrogen and stored at −80°C.

#### Nuclei Isolation and Lysis

1 ml of Mild Cell Lysis Buffer (10 mM Tris-HCl pH 8.1, 10 mM NaCl, 0.5 NP-40, 2X proteinase inhibitors) was applied to cell pellets from 1000 embryos and rotated at 4°C for 10 minutes. Samples were centrifuged at 1300 X g for 5 minutes at 4°C. Supernatant was discarded and nuclei pellets were resuspended in 1 ml Nuclei Wash Buffer (50 mM Tris-HCl pH 8.0, 100 mM NaCl, 10 mM EDTA, 1% SDS, 2X protease inhibitors) and rotated at room temperature for 10 minutes. Samples were centrifuged at 1300 X g for 5 minutes at 4°C to pellet nuclei.

Supernatant was discarded and nuclei pellets were resuspended in 100 µl of Nuclei Lysis Buffer (50 mM Tris-HCl pH 8.0, 10 mM EDTA, 1% SDS, 2X proteinase inhibitors). Samples were incubated on ice for 10 minutes. 900 µl of IP Dilution Buffer (16.7mM Tris-HCl pH 8.1, 167mM NaCl, 1.2mM EDTA, 0.01% SDS, 1.1% Triton X-100, 2X proteinase inhibitors) was added to samples.

#### Chromatin Sonication

Nuclear lysates were sonicated with a Branson Digital Sonifier with the following settings: 10 second duration (0.9 seconds ON, 0.1 seconds OFF), 30% amplitude. 7 sonication cycles were performed. Samples were placed in an ice bath for at least 1 minute between each sonication cycle. Sonicated samples were centrifuged at 14,000 rpm, 4°C, for 10 minutes to pellet insoluble material. Supernatants were transferred to new tubes. A portion of the sample was set aside to confirm optimal chromatin shearing by agarose gel electrophoresis.

#### Preclear

20 µl of Dynabeads (Invitrogen) were blocked with 0.5mg/ml BSA in PBS. Blocked Dynabeads were subsequently applied to each sonicated sample and rotated for 1 hour at 4°C. Samples were placed on a magnet stand for 1 minute and precleared supernatant was transferred to a new tube. 5% of the sample was removed and stored at −80°C as input. Antibody and fresh 1X protease inhibitors were added to each sample. Samples were rotated overnight at 4°C.

#### Pulldown

Samples were centrifuged at 14,000 rpm, 4°C, for 5 minutes to pellet insoluble material. Supernatants were transferred to new tubes. 50 µl of Dynabeads (Invitrogen) were blocked with BSA 5mg/ml in PBS. Blocked Dynabeads were subsequently applied to each sample and rotated for 6 hours at 4°C.

#### Stringency Washes

All wash buffers were kept ice cold during stringency washes. Samples were washed 8 times with RIPA Buffer (10 mM Tris-HCl pH 7.5, 140 mM NaCl, 1mM EDTA, 0.5 mM EGTA, 1% Triton X-100, 0.1% SDS, 0.1% sodium deoxycholate, 2X protease inhibitors), 2 times with LiCl Buffer (10mM Tris-HCl pH 8.0, 1 mM EDTA, 250 mM LiCl, 0.5% NP-40, 0.5% sodium deoxycholate, 2X protease inhibitors), 2 times with TE Buffer (10mM Tris-HCl pH 8.0, 1 mM EDTA, 2X protease inhibitors).

#### Elution and Reversing Crosslinks

100 µl of Elution Buffer (10 mM Tris-HCl pH8.0, 5 mM EDTA, 300 mM NaCl, 0.1% SDS) was added to beads. 2ul of RNase A (Thermo Fisher, Cat #EN531) was added to each ChIP and input sample and incubated at 37°C for 30 minutes with gentle agitation. 10 µl of Proteinase K (Thermo Fisher, Cat # 25530049) was added to each sample and incubated at 37°C for 1 hour with gentle agitation. Crosslinks were reversed overnight at 65°C with gentle agitation. ChIP DNA was purified with a Qiagen MinElute PCR Purification kit (Cat #28004).

#### ChIP-seq Library Construction & Sequencing

ChIP seq libraries were prepared using NEBNext ChIP-Seq Library Prep Reagent Set for Illumina (New England BioLabs, Cat # E6240). High throughput sequencing was performed on Illumina HiSeq 2500 for single-end 50 bp reads or Illumina NovaSeq 6000 for paired-end 50 bp reads.

#### ChIP-rx-seq in Zebrafish Embryos

ChIP-rx was adapted from (Orlando et al., 2014) for H3K27me3 ChIP in zebrafish embryos treated with DMSO or PRT4165. Crosslinked *Drosophila melanogaster* S2 cells (ATCC Cat# CRL-1963) were spiked into resuspended zebrafish embryo pellets at a ratio of 5:1 (zebrafish cells: S2 cells). ChIP-rx was subsequently performed in the same way as described above.

#### ChIP in Zebrafish Sperm

ChIP in zebrafish sperm was conducted as described (Murphy et al., 2018).

#### qPCR

qPCR was carried out using 2X SsoAdvanced Universal SYBR Green Supermix (Biorad Cat # #1725270) and a Biorad CFX real time thermal cycler.

#### Oligonucleotides

See Table S1 for oligonucleotides used for ChIP-qPCR

**Table S1.**
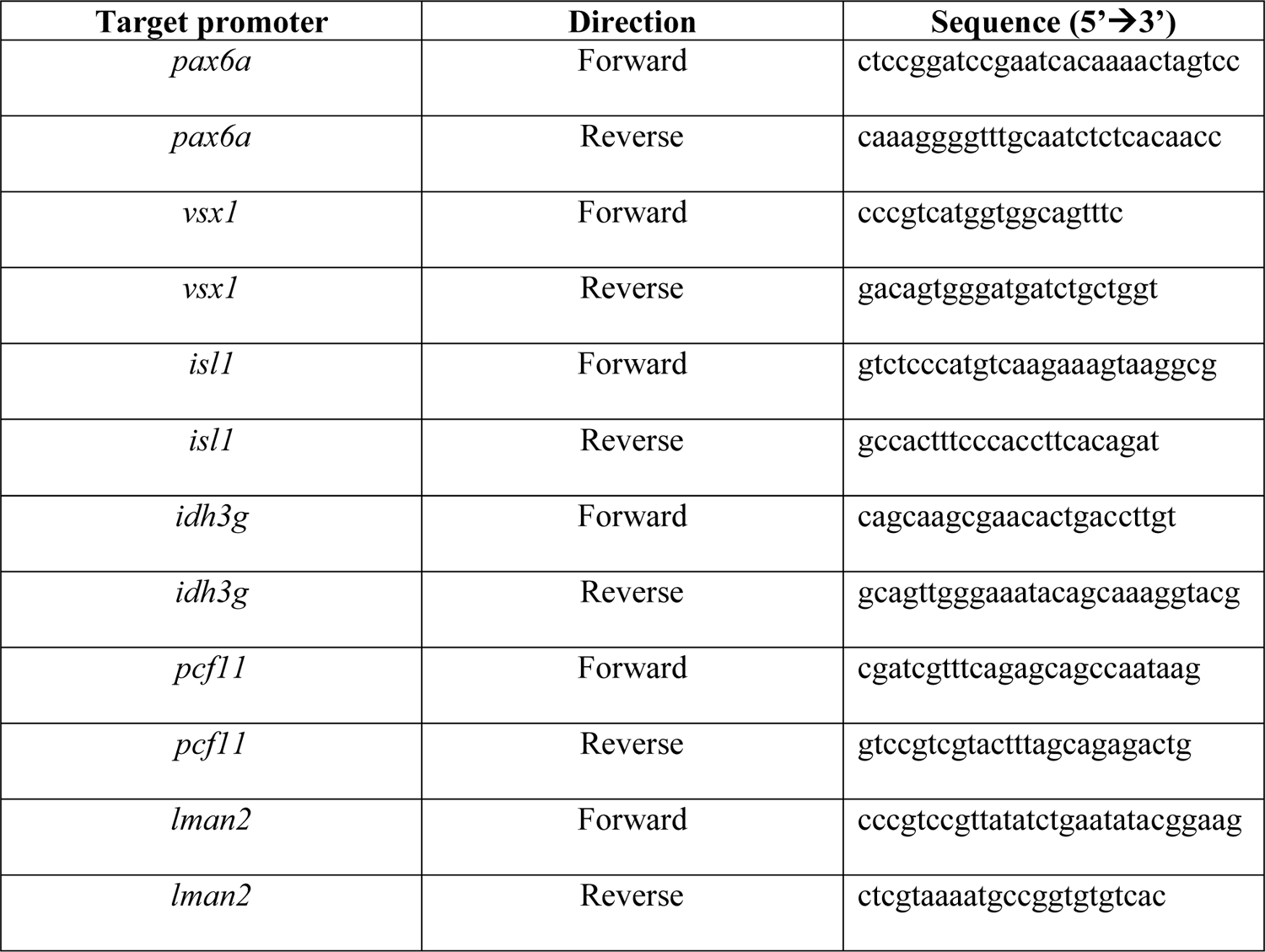
Oligonucleotide sequences used for ChIP-qPCR for amplifying promoter regions.

#### RNA-Seq

Total RNA was harvested from zebrafish embryos with a Qiagen Allprep kit (Cat #80204). The Invitrogen DNA-*free* DNA removal kit (Cat # AM1906) was subsequently used to remove contaminating DNA from RNA samples. Intact poly(A) RNA was purified from total RNA samples (100-500 ng) with oligo(dT) magnetic beads and stranded mRNA sequencing libraries were prepared as described using the Illumina TruSeq Stranded mRNA Library Preparation Kit (RS-122-2101, RS-122-2102). Purified library quality was assessed on an Agilent Technologies 2200 TapeStation using a D1000 ScreenTape assay (Cat # 5067-5582 and 5067-5583). The molarity of adapter-modified molecules was defined by quantitative PCR using the Kapa Biosystems Kapa Library Quant Kit (Cat # KK4824). Individual libraries were normalized to 5 nM and equal volumes were pooled in preparation for Illumina sequence analysis. High throughput sequencing for RNAseq was performed on an Illumina HiSeq 2500. RNA-seq data displayed in Figure 3 – figure supplement 1A was collected from http://www.ebi.ac.uk/gxa/experiments/E-ERAD-475. We would like to thank the Busch-Nentwich lab for providing RNA-seq data used in Figure 3 – figure supplement 1A.

#### ChIP-seq Analysis

ChIP-seq Fastq files were aligned to Zv10 using Novocraft Novoalign with the following settings: -o SAM -r Random. SAM files were processed to BAM format, sorted, and indexed using Samtools (Li et al., 2009). ChIP-seq replica correlation was assessed with deepTools (Ramírez et al., 2016). Briefly, BAM files were read normalized with deeptools bamCoverage with the --normalizeUsingRPKM flag. Deeptools multiBigwigSummary bins, and plotCorrelation were used to generate genome-wide correlation matrices for assessing replica correlation. ChIP-seq peak calling was accomplished using MACS2 with the following settings: callpeak -g 1.4e9 -B -q 0.01 -SPMR (Zhang et al., 2008). Called peaks were annotated in R with the ChIPseeker package (Team, 2020; Yu et al., 2015). Output bedgraph files from MACS2 were processed into bigwig files with UCSC Exe Utilities bedGraphToBigWig (Kent et al., 2010).

Resulting bigwig files were loaded into IGV for genome browser snapshots of ChIP-seq enrichment (Thorvaldsdóttir et al., 2012). Heatmaps of ChIP-seq enrichment at promoter and enhancer regions were made with deepTools (Ramírez et al., 2016). A bed file of Zebrafish zv10 UCSC RefSeq genes from the UCSC Table Browser was utilized for plotting heat maps of ChIP enrichment at promoters (Karolchik et al., 2004). Genes residing on unmapped chromosomal contigs were excluded. Enhancer heatmaps utilized a bed file of postZGA H3K4me1 ChIP-seq (Bogdanovic et al., 2012) peak summits that had been filtered to exclude promoters and unmapped chromosomal contigs. ChIP-seq peaks were annotated by using the Bioconductor package ‘ChIPseeker’ in R (Yu et al., 2015). Gene ontology analysis was performed with DAVID (Huang da et al., 2009).

#### Violin plot of ChIP-seq enrichment at promoters

Log2(ChIP/input) data for promoter regions (+/-1Kb from TSS) of interest was collected from processed bigwig files by utilizing the program Bio-ToolBox ‘get_datasets.pl’ (https://metacpan.org/pod/distribution/Bio-ToolBox/scripts/get_datasets.pl). Collected data was plotted in violin format. Unpaired t-tests with Welch’s correction were utilized to determine statistical differences in ChIP enrichment between promoter K-means clusters. Violin plot and statistical analysis (Figure 1 – figure supplement 2J) were performed in GraphPad Prism version 8.3.1. using GraphPad Prism version 8.3.1 for MacOS, GraphPad Software, San Diego, California USA, www.graphpad.com.

#### DNA Motif Analysis

HOMER was utilized for identifying putative transcription factor binding motifs present at promoters and enhancers (Heinz et al., 2010). The following parameters were used on bed files of promoters and enhancers of interest: findMotifsGenome.pl danRer10 -size −250,250. Known Motifs (as opposed to *de novo* motifs) from HOMER were presented in figures 1 & 2.

#### ChIP-seq analysis involving drug treatments

Analysis of ChIP-seq experiments involving drug treatments was performed by utilizing the Multi-Replica Macs ChIPSeq Wrapper (https://github.com/HuntsmanCancerInstitute/MultiRepMacsChIPSeq).

#### RNA-seq analysis

RNA-seq fastq files were aligned to Zv10 using STAR (Dobin et al., 2012) with the following settings: --runMode alignReads --twopassMode Basic --alignIntronMax 50000 --outSAMtype BAM SortedByCoordinate --outWigType bedGraph --outWigStrand Unstranded -- clip3pAdapterSeq AGATCGGAAGAGCACACGTCTGAACTCCAGTCA. The resulting sorted BAM files were subsequently indexed using Samtools (Li et al., 2009). FeatureCounts was utilized to collect count data for zv10 genes via the following command: -T 16 -s 2 – largestOverlap (Liao et al., 2013). Count data for all replicates across experimental conditions were combined into a single count matrix in R (Team, 2020). This count matrix was subsequently used to identify differentially expressed genes with the R package DESeq (Anders et al., 2010). RNA-seq replica correlation was assessed with deepTools (Ramírez et al., 2016).

Briefly, BAM files were read normalized with deeptools bamCoverage with the -- normalizeUsingRPKM flag (Ramírez et al., 2016). Deeptools multiBigwigSummary bins, and plotCorrelation were used to generate genome-wide correlation matrices for assessing replica correlation (Ramírez et al., 2016).

#### Reprocessed ChIP-seq Datasets

PreZGA H2Az & preZGA H3K4me1 ChIP-seq data (Murphy et al., 2018) (GEO: GSE95033), postZGA H3K4me1 ChIP-seq data (Bogdanovic et al., 2012) (GEO: GSE68087), preZGA & postZGA H3K27ac ChIP-seq data (Zhang et al., 2018) (GEO: GSE114954), postZGA H3K4me3 & H3K27me3 ChIP-seq data (Zhang et al., 2014) (GEO: GSE44269) and Nanog ChIP-seq (Xu et al., 2012) (GEO: GSE34683) were downloaded from the Gene Expression Omibus and reprocessed as described above.

#### Whole Genome Bisulfite Sequencing Analysis

WGBS from Potok et al., 2013 (DRA/SRA: SRP020008) was processed as described (Murphy et al., 2018).

#### Data Access

All sequencing datasets generated in this study have been deposited at the Gene Expression Omnibus under the accession number GSE168362.

#### Drug Treatments

PRT4165 (Tocris Cat #5047) was dissolved in DMSO at a concentration of 50 mM. PRT4165 was further diluted to a working concentration of 150 µM in embryo water and mixed vigorously. Zebrafish embryos were collected and immediately placed in embryo water containing 150 µM PRT4165 (or DMSO) and allowed to develop to 4 hpf.

## Acknowledgements

We thank Brian Dalley and the Huntsman Cancer Institute High-Throughput Sequencing Shared Resource for sequencing data. We thank Tim Parnell, Chongil Yi, and Jingtao Guo for helpful discussions regarding bioinformatic analysis. We thank David Grunwald and Rodney Stewart for providing helpful feedback during manuscript preparation. We thank all current and former members of the Cairns lab for their perspectives on this project. B.R.C. is an investigator with the Howard Hughes Medical Institute. C.W. was funded by the T32 Developmental Biology Training Grant (59202072), 4DNucleome (NIH Common Fund) NBR - 92275293 S9001779, and HFSP RGP0025/2015. Imaging was performed at the University of Utah Microscopy Core (1S10RR024761-01). Financial support was received from the Howard Hughes Medical Institute and the Huntsman Cancer Institute core facilities (CA24014).

